# Predicting the impact of giant kelp restoration on food webs and fisheries production

**DOI:** 10.1101/2024.12.16.628810

**Authors:** Tess O’Neill, Scott Bennett, Christopher J. Brown

## Abstract

1. Ecosystem restoration is gaining momentum as communities and policymakers increasingly appreciate the need to recover lost ecosystem services. However, more knowledge about the potential ecological outcomes of restoration for ecosystem services is needed to promote engagement of stakeholders with restoration targets. In Tasmania the giant kelp, *Macrocystis pyrifera,* is a productive canopy species that has declined by 95%.

2. We aimed to predict the effects of *M. pyrifera* restoration on foodwebs and fisheries production. The primary productivity multiple of *M. pyrifera* compared to the dominant understory species *Ecklonia radiata* was quantified and then incorporated into an Ecopath with Ecosim (EwE) model of Tasmanian waters. The model was fitted to catch per unit effort data to estimate the predator/prey interactions. Several scenarios were then simulated, representing uncertainty in predator/prey interactions and different areas of *M. pyrifera* restoration.

3. Restoration of degraded reefs with *M. pyrifera* was predicted to increase primary productivity by about 40 times per unit area compared to the existing habitat.

4. We predicted that restoring 30% of the degraded *M. pyrifera* area would increase abalone and coastal demersal fish catch rates by ∼7%. Rock lobster and reef-associated fisheries catch rates were predicted to increase by 1-2% respectively.

5. Model scale was hypothesized to underestimate the increases in biomass and catch. By modelling outcomes of ecological restoration, achievable targets can be set that are locally relevant and therefore more likely to attract support.

## 1. Introduction

Coastal habitat restoration is increasing globally to aid the recovery of degraded ecosystems (Basconi et al., 2020; Saunders et al., 2020). Coastal habitat restoration is usually defined as assisted regeneration of coastal ecosystems and can involve transplanting or seeding habitat-forming species, improving environmental conditions to promote natural recovery and removing grazers (Gann et al., 2019). Recovery of coastal habitats is sought after by communities and governments because it contributes to conservation objectives (Possingham et al., 2015) and can enhance ecosystem services including carbon sequestration, coastal protection, nutrient filtration and fisheries (Barbier et al., 2011; Unsworth et al., 2019). Restoration is forecast to soon be at a scale that benefits biodiversity and ecosystem services across seascapes (Saunders et al., 2024).

Global policies and organisations recognise the need for ecosystem restoration and the intended outcomes of enhancing biodiversity and ecosystem services. For example, Target 2 of the Kunming-Montreal Global Biodiversity Framework (KMGBF) states: “Ensure that by 2030 at least 30% of areas of degraded…coastal ecosystems are under effective restoration…to enhance biodiversity and ecosystem functions and services…” (Convention on Biological Diversity, 2022). Global targets are typically articulated in terms of areas of habitat restored, but lack of tangible outcomes for ecosystems and people can hinder motivation towards restoration efforts (Bell-James et al., 2024). Therefore, it is necessary to align global restoration targets with their outcomes for biodiversity and ecosystem services. Predictions for outcomes, particularly those with local relevance, can help clarify the expectations communities have about restoration efforts and improve engagement from stakeholders who are seeking ecosystem services, such as fisheries or carbon sequestration, from restoration (Bayraktarov et al., 2020; Saunders et al., 2024).

Coastal habitat restoration can improve biodiversity and ecosystem services, but this is determined by how animals respond to habitat recovery. Such responses are variable and often based on short-term datasets that may not represent the long-term impacts of successful restoration (Sievers et al., 2024). Linking habitat restoration to ecosystem change and fisheries production requires knowledge about the use of the coastal habitats by animals and complex food web dynamics (Basconi et al., 2020; Sievers et al., 2022). Therefore, with the limited number of successful examples to provide empirical evidence (Saunders et al., 2020), ecosystem models provide a useful tool to predict the impact coastal habitat restoration on food webs and secondary production (Vasslides et al., 2017). Food web models, such as Ecopath with Ecosim (Ewe; Christensen & Walters, 2004), can help resolve the flows of primary production from restored habitats through a food web and the resulting biomass dynamics of secondary producers (Heymans et al., 2016; Geary et al., 2020). Food web models need to consider uncertainty about predatory-prey interactions, which influence the response to habitat changes (Christensen & Walters, 2004). Although primarily used to predict ecological impacts of climate change and fisheries policies (Geary et al., 2020), food web models are increasingly used to estimate the outcomes of coastal habitat restoration (Vasslides et al., 2017) and have predicted enhancement of animal biomass following the restoration of coastal habitats, such as seagrass and oyster reefs (Horn et al., 2021).

Here, we apply food-web modelling to simulate the ecosystem and fisheries outcomes of Giant Kelp (*Macrocystis pyrifera*) restoration in Tasmania, Australia. *M. pyrifera* forests in Tasmania have declined by 95% over recent decades in association with ocean-warming and an intensification of the East Australian Current that carries warm, nutrient poor water to eastern Tasmania (Johnson et al., 2011; Butler et al., 2020). These severe declines prompted the listing of *M. pyrifera* forest communities in southeastern Australia as ‘endangered’ by the Australian Government in 2012 (DCCEEW, 2022). Restoration efforts are underway to promote the recovery of *M. pyrifera* forest area towards global targets (Eger et al., 2024; Layton et al., 2020). Kelp is well recognised as a key habitat-forming species that supports a range of animals (Schiel & Foster, 2015; Steneck et al., 2002), however the influence of kelp productivity on structuring kelp forest food webs is less clear (Elliott Smith & Fox, 2022; Hobbs, 2007). Estimates of kelp productivity are also highly variable and often summarized at global scales or broad taxonomic levels (Pessarrodona et al., 2022). *M. pyrifera* is one of the fastest growing kelp species, and its recovery could significantly enhance primary productivity of a seascape, with flow on effects to secondary production. Quantifying the impact of restoring *M. pyrifera* on primary productivity requires site-specific estimates of *M. pyrifera* productivity versus the species it replaces, generally the less productive kelp species, *Ecklonia radiata*.

To address these knowledge gaps, we simulated *M. pyrifera* restoration within a Tasmanian EwE food web model (Watson et al., 2013) to predict changes in secondary production with a focus on important fisheries groups. This study is also intended as an informative case study for using ecosystem food web models to translate area-based restoration goals into outcome-related targets that can be better understood by communities and increase motivation for stakeholder engagement. We aimed to answer the following questions: 1) what is the productivity of *M. pyrifera* compared to *E. radiata* in Tasmania 2) does updated fisheries data parameterise predator-prey interactions differently than initial fitting by Watson et al. (2013) and 3) what are the predicted changes in biomass of consumers within the Tasmanian ecosystem following *M. pyrifera* restoration?

## 2. Methods

### 2.1. Overview of approach

There were 3 main steps: 1) Estimate the primary productivity ratio of *M. pyrifera* compared to *E. radiata*, to represent the enhanced productivity per unit area from *M. pyrifera* restoration 2) parameterise predator-prey interactions by fitting the EwE to fisheries data; and 3) applying the average productivity ratio as a ‘productivity multiplier’ to simulate different *M. pyrifera* restoration scenarios in the model and predict changes to relative biomass and fishery catch for animal functional groups.

### 2.2. Productivity of *M. pyrifera* compared to *E. radiata*

To represent the productivity changes from replacing *E. radiata* with *M. pyrifera* through restoration, we matched the productivity of both species for the same seasons and sites with comparable environmental conditions and calculated productivity ratios of *M. pyrifera* compared to *E. radiata* for each matched sample.

#### 2.2.1. Summary of methods for productivity data

The biomass accumulation values, and density estimates used in this study were collected by Bennett et al. (in review) during surveys conducted across six study sites multiple times from 2021-2023. *M. pyrifera* growth rates were measured at three ‘*M. pyrifera* forest’ sites selected for their large areas of tall, dense canopies. Growth rates of *E. radiata* were also measured where this species was present at these sites as well as the three additional ‘*E. radiata* forest’ sites where *E. radiata* was the dominant species and *M. pyrifera* was not present. The productivity metric calculated for both species was a measure of biomass accumulation over time i.e. the increase in wet weight (g) individual frond^-1^ day^-1^. The methods used were appropriate to the different growth forms (Sanderson, 1990; Schiel & Foster, 2015). Density estimates were obtained from quadrat surveys of *M. pyrifera* and some *E. radiata* forests.

During different seasons throughout 2021-2023, mature individuals of *E. radiata* at the six sites were tagged and hole-punched along the lamina at known distances from the stipe-meristem junction and the top of the lamina. Throughout the same time period, *M. pyrifera* stipes of different individuals at the three *M. pyrifera* forest sites were tagged with tape 1m from the tip (Sanderson, 1990; Schiel & Foster, 2015). The number of blades between the tip and the tape was counted as well as the total number of stipes on the *M. pyrifera* individual. After one month, all tagged individuals were harvested and measured in the laboratory.

For *E. radiata*, the distances between the punched holes were re-measured and compared to the original distances at the time of tagging. Total weight, lamina length and weight per section of the individual were measured. Calculations using these measurements consider growth and erosion to ultimately calculate the daily growth of the individual. For *M. pyrifera*, distance and number of blades between the tape and tip were re-counted and the total weight and length was measured. The stipe was cut 1m above the tape and the remaining segment of the tip was weighed as a measure of the new biomass accumulated during the survey period. The growth of the whole *M. pyrifera* individual was calculated by multiplying the growth of the stipe by the total number of stipes on the individual and resulted in a measurement of daily growth per individual.

#### 2.2.2. Ratio of *M. pyrifera* to *E. radiata* productivity

We grouped and averaged productivity values of *E. radiata* by location, site, year and season and assigned unique names to the samples. Density values of the number of individuals of *E. radiata* per quadrat were converted to individuals per 5m^2^ and filtered to contain only adult *E. radiata* fronds. The density data was then matched to the productivity samples. In cases where there was not a direct match, the density value of a similar sample was matched with multiple productivity samples that had similar sites, because *E. radiata* density remains relatively stable between years and seasons (S. Bennett, personal observation, 2024). For each sample, the average productivity per 5m^2^ area was calculated by multiplying the average productivity per individual by the number of individuals per 5m^2^. We used the same steps to estimate productivity values of *M. pyrifera* productivity per 5m2 for each sample.

There were 20 unique site x season combinations for *E. radiata* and nine unique site x season combinations for *M. pyrifera* productivity in g per 5m^2^. Samples of *E. radiata* and *M. pyrifera* were matched to calculate productivity ratios that represented the replacement of *E. radiata* by *M. pyrifera* beds at the site scale. Five of these were exact matches, and the remaining *E. radiata* samples were matched with *M. pyrifera* samples that had similar environments (i.e. adjacent sites within ∼1nm). Productivity varies between year and season for both kelp species (S. Bennett, personal observation, 2024) so these variables were kept consistent between pairs. Due to the proximity between Actaeon and Southport sites, and their similar forest dynamics such as light intensity, Southport sites and Actaeon sites were matched in cases where *E. radiata* was missing from Acteon surveys.

Productivity ratios for each sample were calculated by dividing *M. pyrifera* productivity by *E. radiata* productivity. These ratios represented the net productivity benefit per unit area if *M. pyrifera* replaced *E. radiata*.

### 2.3. Ecological modelling

#### 2.3.1. Tasmanian Waters EwE model

We used an EwE model that represented the marine shelf environment surrounding Tasmania (Watson et al., 2013). The model represented nearly 137,000km^2^ and includes 47 functional groups and 16 fisheries. The general structure and approach of EwE has been described and reviewed in detail (Christensen & Walters, 2004; Heymans et al., 2016) so is briefly summarised here. Ecopath is the mass-balance routine that represents a food-web and describes the ecosystem using ‘functional groups’, a species or group of species that share a similar habitat, diet or overall function within the system. Functional groups can be assigned as either producers, consumers or detritus. The biomass, productivity, consumption and ecotrophic efficiency of groups are based on information from empirical studies and fisheries data, and inbuilt Ecopath equations balance the model to estimate missing information.

Ecosim is the time dynamic module of the EwE software that allows users to predict dynamic changes in biomass. Ecosim can be used to predict the responses of biomasses to perturbations in the system, such as changes to fishing mortality or primary production (Christensen & Walters, 2004). Biomasses change as a function of production and losses and are determined by a master equation for each functional group *i*:

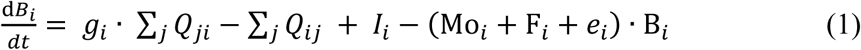

where *B* is the biomass, *g* is the growth efficiency, *Q* is the consumption rates of all prey *j* by group *i* (*Cji*) and of group *i* by all predators *j* (*Cij*), *I* is the immigration rate, Mo is the mortality not attributable to other model groups, *F* is the fishing mortality rate, and *e* is the emigration rate.

The consumption rate is defined as

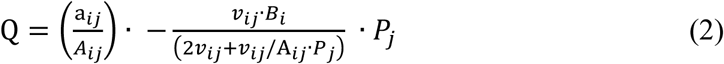

where *aij* is the rate of effective search for *i* by *j*, *Aij* is the search area in which *j* forages for *i*, *vij* is the flow rate of *Bi* between pools that are invulnerable and vulnerable to predator *j*, and *Pj* is the abundance of *j* in *Aij*.

Trophic interactions drive the consumption rates within the system and are based on the foraging arena theory (Ahrens et al., 2012): prey biomass pools are dynamically divided into components that are available and unavailable to predation (Bentley et al., 2024; Christensen & Walters, 2004). The rate of transfer of a prey group between these two states is controlled by the ‘vulnerability’ parameter, which can shift predator-prey interactions within the system. Ecosim predictions are highly sensitive to the vulnerability parameter.

#### 2.3.2. Vulnerability fitting

Vulnerability settings were calibrated in the model through a fitting procedure. The fitting process in Ecosim involves using forcing data to drive changes in the model and find evidence-based vulnerability parameters by minimising the sums of squares between hindcasts of relative biomass from Ecosim with an observed timeseries of relative biomass (often catch per unit effort (CPUE) data). The original model parameterised vulnerabilities by fitting the model to various Tasmanian fisheries CPUE timeseries from 1995-2007 (Zeigler and Lyle, 2009 in Watson 2013) and here we fit the model using updated fisheries data from 1995 to 2022, including adding effort and CPUE data of *Haliotis rubra* (Blacklip Abalone), which was not included in the initial fitting. Abalone is of significant interest due to kelp forests being its primary habitat and its high economic value for Tasmania.

Fitting was completed in two stages (Christensen and Walters 2004). First, we applied one forcing function to the two phytoplankton groups and a second to benthic primary producers (macroalgae and seagrass) and used the primary productivity anomaly search capabilities to identify trends that matched the fishing data. We limited it to two forcing functions to prevent overfitting. The primary productivity anomaly search on both forcing functions was ran using 9 knots in the spline. This allowed the primary productivity forcing functions to represent broad productivity trends throughout the time period of available fisheries data but without overfitting by allowing year to year variation. We then ran the vulnerability sensitivity search to identify the most sensitive vulnerabilities by predators.

Finally, we adjusted the top 10 most sensitive predator vulnerabilities to fit to the fisheries data to obtain our final vulnerability matrix. We compared the AIC with searches for 5 and twenty predators finding that 10 predators had the lowest AIC and therefore the most parsimony. Fitted vulnerabilities were compared to the values initially parameterised by Watson et al. (2013).

#### 2.3.2. *M. pyrifera* restoration scenarios

A series of *M. pyrifera* restoration scenarios were simulated in Ecosim with each scenario representing different restoration areas and vulnerability settings. The EwE model contained a ‘Macroalgae’ functional group (Watson et al., 2013), which represents the dominant canopy forming kelps, primarily E. radiata. *M. pyrifera* restoration was represented as a one-off, step increase in macroalgae productivity that persisted through time until the model groups’ stabilised. Three *M. pyrifera* restoration areas were simulated: 1) 0.7ha has recently been restored at a trial site in Tasmania; 2) 10ha is the short-term target for Tasmania based on funding and 3) the ‘30 by 30’ KMGBF target. Each scenario had a vulnerability setting representing a bottom-up system (vulnerability = 2.0), top-down system (vulnerability = 5.0), vulnerabilities fitted to fisheries data from the original study (Watson et al., 2013) or vulnerabilities fitted to updated fisheries data in this study.

To simulate the global biodiversity restoration target in the model, we converted the ‘30%’ into an area of *M. pyrifera* restoration. The target calls for 30% of degraded ecosystems to be under effective restoration and therefore requires knowledge about how much *M. pyrifera* has been degraded, or in Tasmania’s case, lost almost entirely. In consideration of the statewide scale of the EwE model, the statewide historical *M. pyrifera* coverage estimate of 422ha in 1989 was used along with the estimate that 95% of *M. pyrifera* has been lost (Butler et al., 2020; Johnson et al., 2011). This is likely a conservative estimate of coverage based on a baseline that was already in decline and previous estimates predict ∼1000ha of *M. pyrifera* coverage at individual regions of Tasmania alone (Johnson et al., 2011).

We calculated the area of *M. pyrifera* restoration that the KMGBF restoration target calls for as:

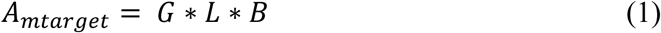

Where *G* is the proportion that the KMGBF target calls for degraded ecosystems to be under effective restoration, *L* is the proportion that *M. pyrifera* is estimated to have been lost in Tasmania and *B* is the statewide historical estimate of *M. pyrifera* coverage.

We determined the productivity forcing function associated with different areas of *M. pyrifera* restoration with the assumption that the initial productivity in the macroalgae group represented *E. radiata* only. We calculated the productivity forcing function in a year *t* as:

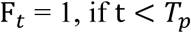

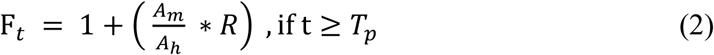

Where *T_p_* was the year restoration started, set to four years so that the model stabilised. *A_m_* was the area of *M. pyrifera* restoration for a given scenario, *A*_ℎ_ was the total reef area in the model and *R* was the *M. pyrifera* to *E. radiata* productivity ratio.

Changes in relative biomass and catch were determined by the Ecosim output after the biomass’ stabilised following the forcing function step increase and were averaged over 50 years in scenarios where relative biomass’ reached a dynamic equilibrium, usually in scenarios with high vulnerability settings.

## 3. Results

### 3.1 Productivity ratio of *M. pyrifera* to *E. radiata*

Average productivity ratios for *M. pyrifera* and *E. radiata* were produced for twenty unique sample names (Figure 1). The average *M. pyrifera* productivity in these samples ranged from 111.17 – 969.3 g 5m^2-1^ day^-1^ and the average *E. radiata* ranged from 2.76 g 5m^2-1^ day^-1^ to 43.03 g 5m^2-1^ day^-1^. The mean productivity for *M. pyrifera* and *E. radiata* was 603.37 g 5m^2-1^ day^-1^ and 19.09 g 5m^2-1^ day^-1^ respectively. The median *M. pyrifera*:*E. radiata* ratio was 40.5, and ranged from 9 to 90. Notably, *M. pyrifera* productivity in each matched sample was always greater than *E. radiata* productivity.

**Figure 1.**
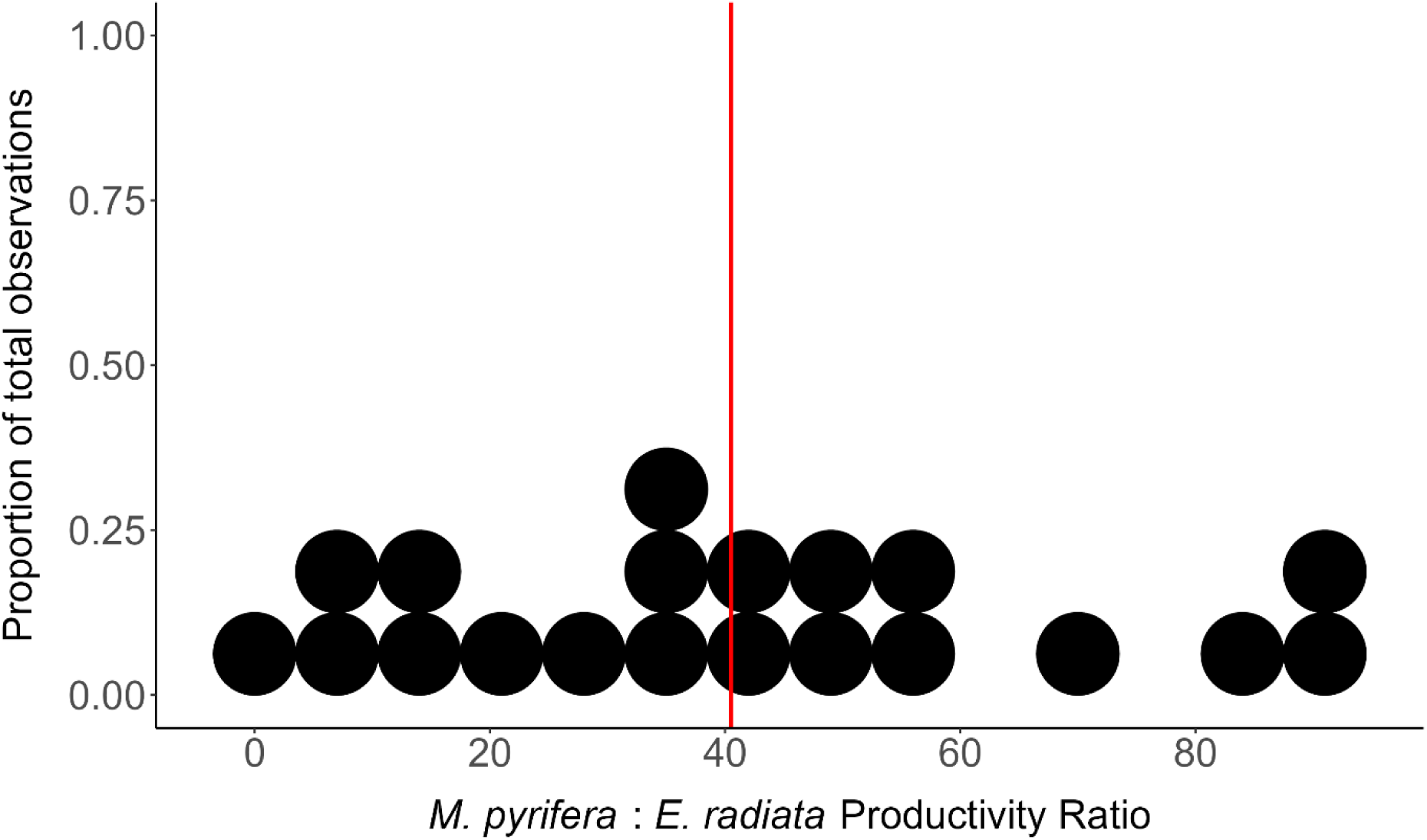
Productivity ratios of *M. pyrifera* compared to *E. radiata* from 20 samples. Dots represent each sample. The red line represents the median ratio (40.5).

### 3.2. Vulnerability fitting

The model was fitted to observed CPUE trends for rock lobster, southern calamari, bastard trumpeter, southern garfish and abalone (Figure 2). The base model fitted biomasses with a sums of squares = 5.133 and an Akaike Information Criteria = -409.8, compared to 9.3 before fitting.

**Figure 2.**
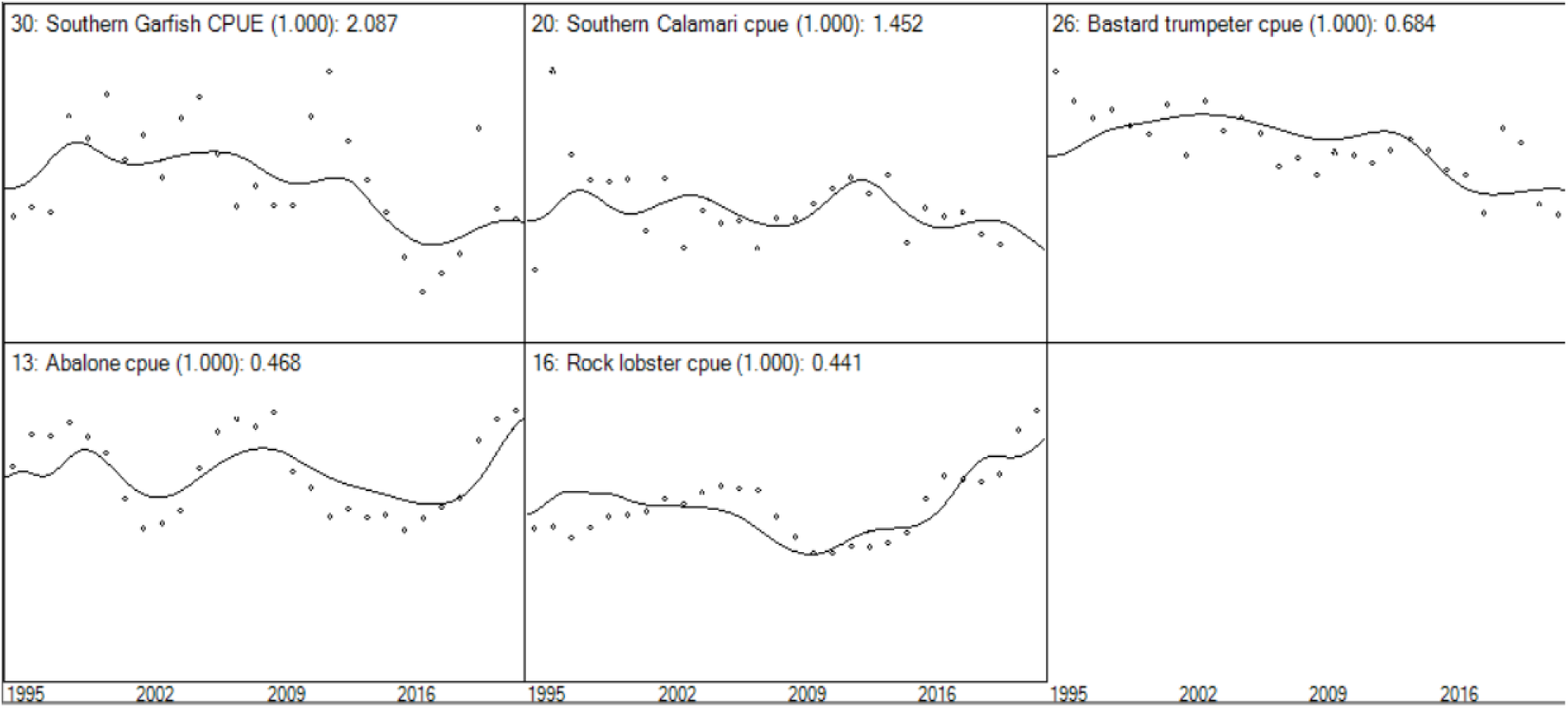
Observed (dots) and predicted (solid lines) biomass for the model for functional groups 30, 20, 26, 13 and 16. Note that many functional groups include more than one species and CPUE data from the species specified here was used to represent the entire group. Values following colon in titles refer to sums of squares for each group.

### 3.3 *M. pyrifera* restoration primary productivity forcing functions

The *M. pyrifera* restoration area for the KMGBF target was estimated to be 120ha which transferred into a 1.8% in macroalgae productivity. The macroalgae productivity multipliers for 10ha and 0.7ha were 0.15% and 0.01% respectively.

### 3.4. *M. pyrifera* restoration simulations

#### 3.4.1. KMBGF target *M. pyrifera* restoration scenario

The changes in relative biomass of functional groups in response to 120ha of *M. pyrifera* restoration could be categorized into three groups: 1) Increases in relative biomass, 2) Small increases or decreases that were likely inconsequential to the system at a statewide scale but still highlighted predicted direction of change in biomass and 3) No change in biomass relative to the initial balanced model (Figure 3).

**Figure 3.**
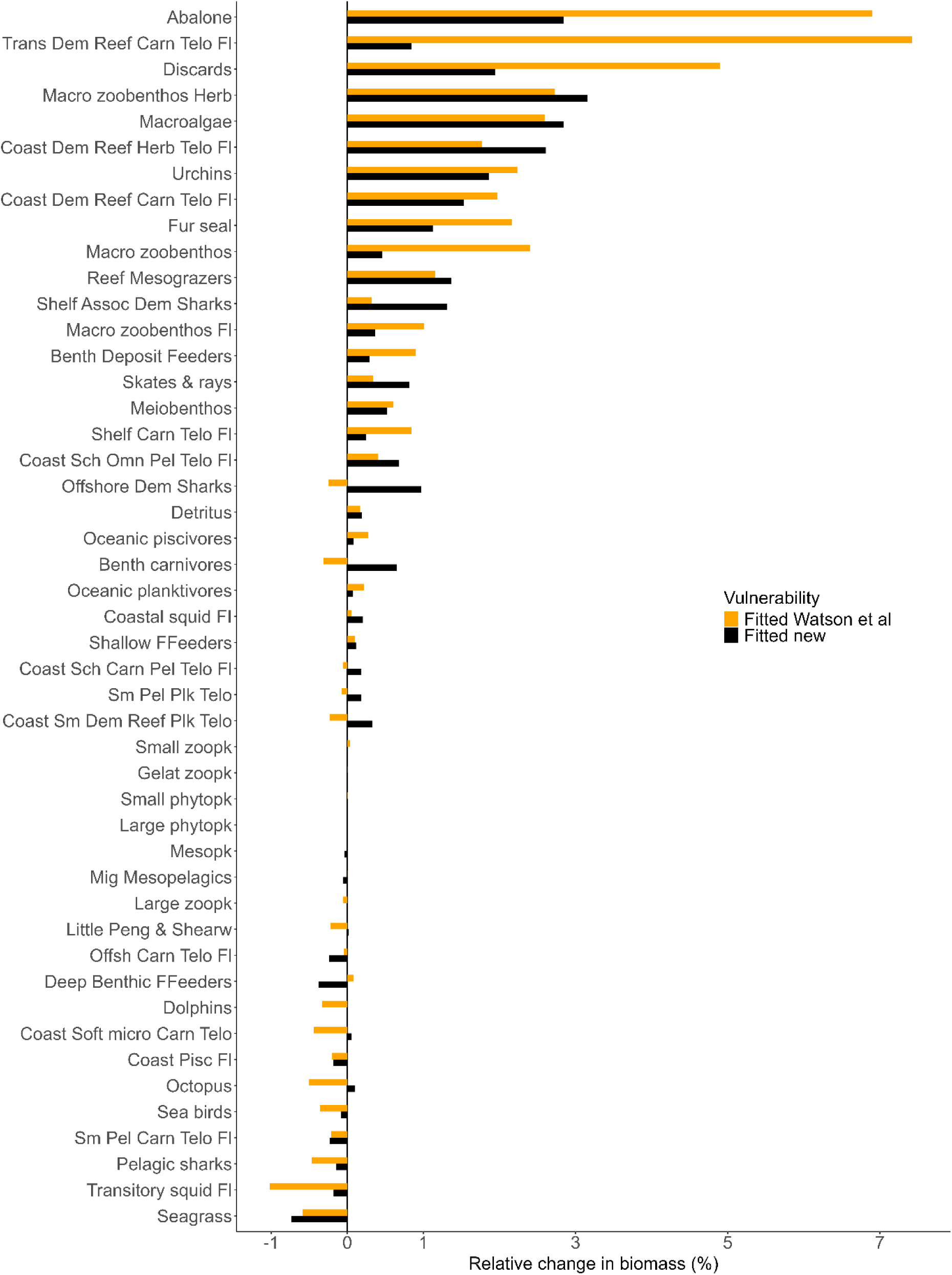
Changes in biomass of model groups after simulating 120ha *M. pyrifera* restoration in Ecosim, relative to initial Ecopath values (biomass’ stabilised ∼70 years after restoration). Macroalgae productivity was increased by 1.8%. Vulnerabilities were parameterised by Watson et al. (2013; yellow) and fitting procedures performed in this study (black).

The relative biomass increases to coastal demersal reef herbivorous teleosts in Watson et al.’s (2013) fitted models were directly proportionate to the relative increase in macroalgae productivity in each forcing function. For instance, following a 1.8% increase in macroalgae productivity, the relative biomass of coastal demersal reef herbivorous teleosts was predicted to increase 1.8%. All other groups either showed changes in relative biomass greater or less than the increase in macroalgae productivity.

The biomasses of most reef-associated functional groups were predicted to increase under the fitted models’ 1.8% scenario (Figure 3). For example, transitory demersal reef carnivore teleosts and abalone were predicted to increase by 7.4% and 6.9% respectively. Some groups, mostly coastal or pelagic groups, were predicted to decrease in biomass following the restoration but by < 1%. Overall, the biomass of plankton and planktivore functional groups were predicted to undergo little change following *M. pyrifera* restoration.

The direction of predicted changes for functional groups’ biomasses between the two fitted models was similar, with the exception of benthic carnivores, offshore demersal sharks and coastal small demersal planktivores. There were significant discrepancies in the magnitude of predicted change in biomass for many functional groups between the two fitted models. For example, Watson et al’s (2013) fitted model predicted greater increases in biomass of abalone and transitory demersal carnivore teleosts but smaller increases in the biomass of shelf associated demersal sharks and coastal demersal reef herbivore teleosts when compared to the model fitted to updated data.

The fitted models predicted increases in relative catch in 8 out of the 16 fleets following the 120ha restoration scenario with fitted vulnerabilities (Figure 4).

**Figure 4.**
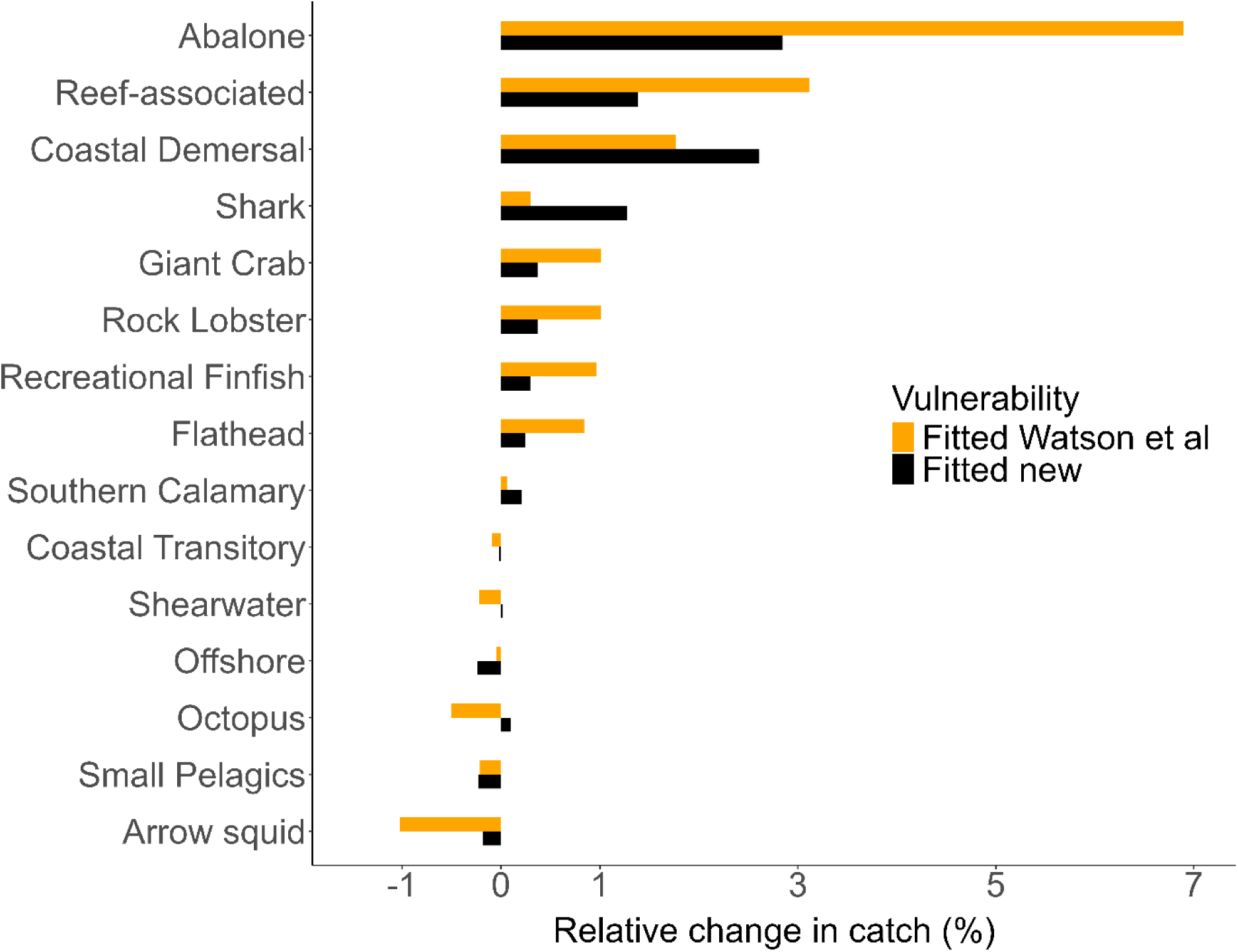
Changes in catch of model groups after simulating 120ha *M. pyrifera* restoration in Ecosim, relative to initial Ecopath values (catches stabilised ∼70 years after restoration).

Macroalgae productivity was increased by 1.8%. Vulnerabilities were parameterised by Watson et al. (2013; yellow) and fitting performed in this study (black). Rock lobster includes both commercial and recreational fleets.

Abalone catch showed the largest relative increase (6.9% in Watson et al.’s (2013) fitted model and 2.8% in updated fitted model), followed by reef-associated and coastal demersal (Figure 4). Other fleets that are economically important for Tasmania showed increased catches but to a lesser extent (∼1%), such as the commercial and recreational rock lobster fleets and the recreational finfish fleet, which included ten functional groups.

EwE uses change in biomass to determine changes in catch and fishing pressure was kept consistent throughout the simulation period. As such, for the fleets that contained only one functional group e.g. abalone, rock lobster, the predicted change in relative catch was equivalent to the predicted change in biomass of those groups (Figure 4). In fleets that contained multiple functional groups e.g. recreational finfish, the catch was disproportionate to the average change in relative biomass of all groups. For example, in the Watson et al. (2013) model, the average increase in biomass within the recreational finfish fleet was 1.8%, however the relative catch increase for the fleet was only 1%. This is due to the inclusion of non-reef associated fauna within the fleet, half of which were predicted to decrease in biomass. Fleets containing reef groups were generally predicted to experience increased catch following the 120ha scenarios in the fitted models whereas the catch of fleets containing plankton consuming groups e.g. coastal transitory fleet, were not predicted to change significantly. Fleets that were predicted to decrease in catch were consistent with their associated functional groups’ predicted changes in biomass.

The recreational finfish fleet includes ten functional groups, only one of which is not included in the other fleets (oceanic piscivores e.g. tuna, swordfish, billfish). Therefore, the increases in recreational finfish fleet are consistent with the increases in other fleets. The giant crab and rock lobster fleets include the same functional group; carnivorous macro zoobenthos fished, and so catch increases were identical.

#### 3.4.2. Effect of vulnerability parameterisation

The predicted changes in relative biomass of functional groups following restoration scenarios differed between the vulnerability parameters used (Figure 5). However, the specific groups that had ecologically insignificant predicted changes in biomass were consistent for all vulnerability settings. Similarly, the groups that were predicted to increase the most were in general the same for all vulnerability parameters. These groups included reef associated groups such as abalone, transitory demersal reef carnivores and macrozoobenthic carnivores and herbivores.

**Figure 5.**
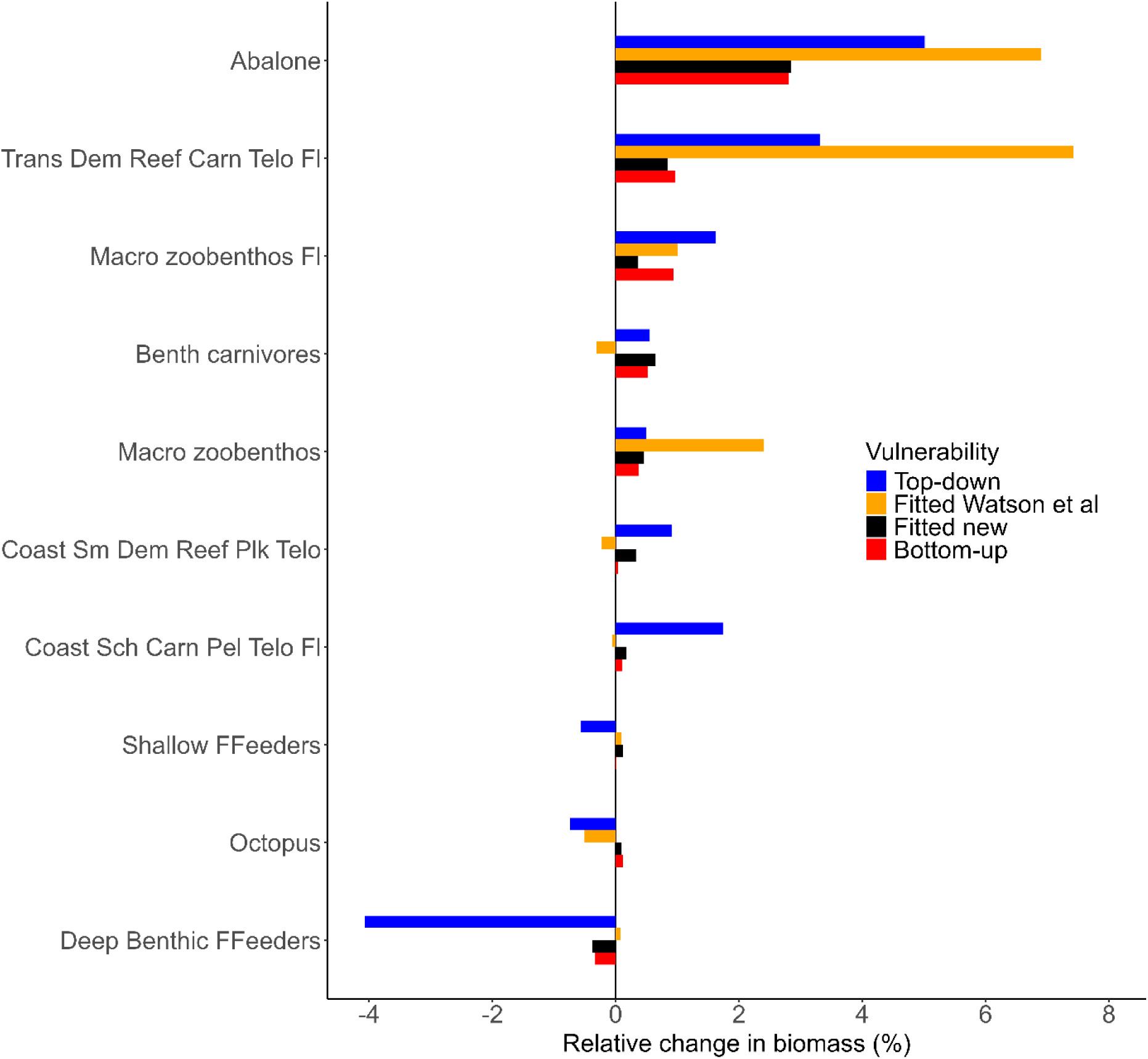
Predicted changes in relative biomass of some functional groups from models with different vulnerability settings following a 120ha *M. pyrifera* restoration scenario.

The top-down model tended to predict larger changes in relative biomass than the bottom-up model. The biomass predictions from the updated fitted model were most similar to the predictions made by the bottom-up model. Differences in biomass change between the fitted and bottom-up models are due to one or other of the fitted models having high vulnerabilities than the other model, and therefore stronger top-down control, for some predator-prey interactions.

#### 3.4.3. Restoration areas

The macroalgae productivity increase under the 120ha *M. pyrifera* restoration scenarios was twelve times greater than that of the 10ha scenarios. In the bottom-up models, the changes in relative biomass and catch were almost proportional to the productivity differences (Figure 6). However, for the fitted and top-down models, the changes in biomass did not align proportionally with the different productivity increases applied (Figure 6).

**Figure 6.**
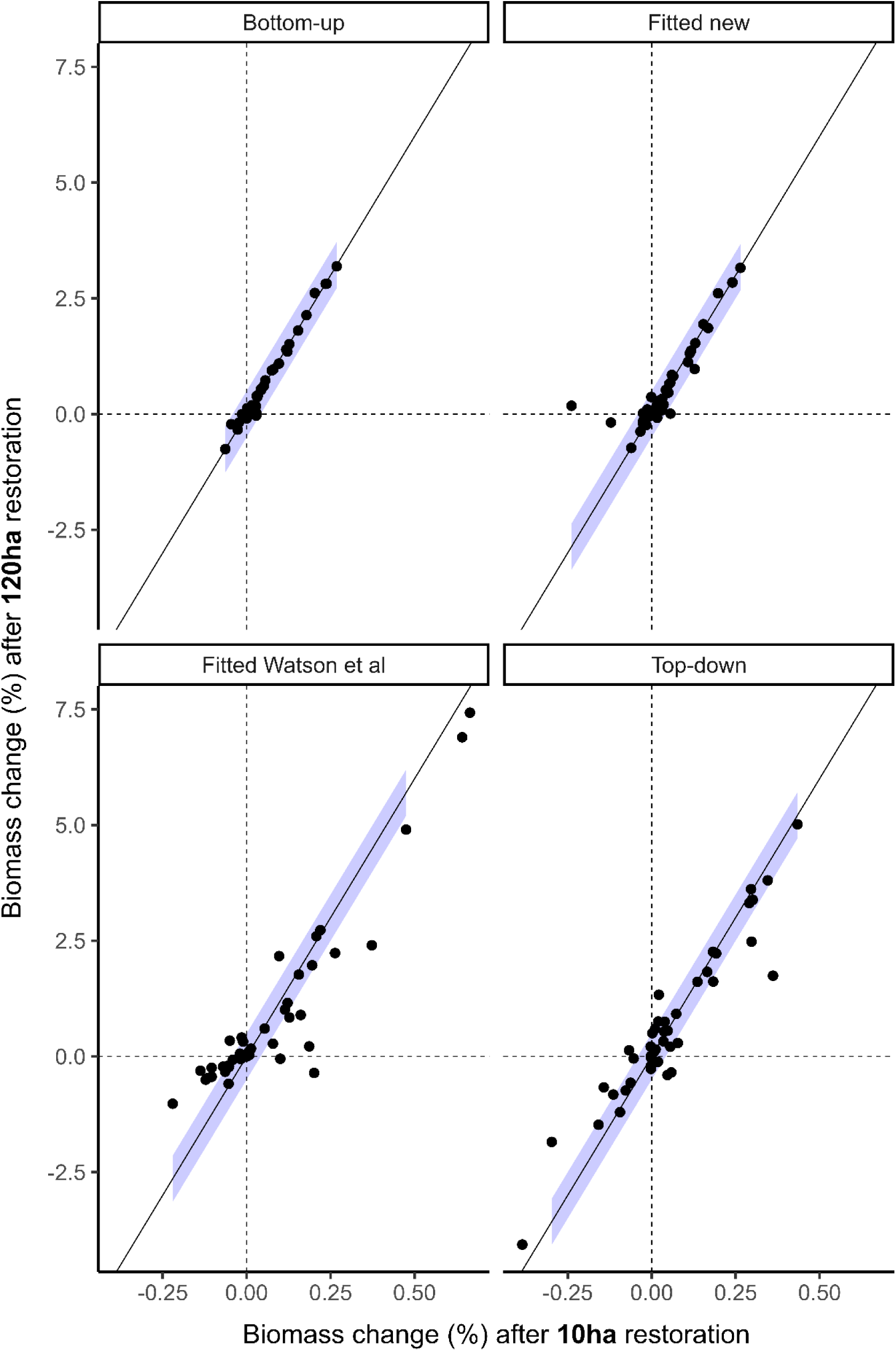
Relative biomass changes of functional groups (points) following 120ha and 10ha of *M. pyrifera* restoration simulated in models with different vulnerability settings. Bottom-up model = all vulnerabilities set to 2.0; Fitted new model = vulnerabilities fitted to updated fisheries data in this study; Fitted Watson et al. (2013) model = vulnerabilities from initial fitting in Watson et al. (2013); Top-down model = all vulnerabilities set to 5.0. Solid line (slope = 12) represents the relative proportion of the 120ha forcing function (1.8%) to the 10ha forcing function (0.15%). Blue shading is +/- 0.5% difference from the expected proportional biomass change.

Specifically, the fitted and top-down models predicted biomass increases for 10ha restoration simulations to be greater than one twelfth of those predicted for the 120ha restoration. For instance, in Watson et al.’s (2013) fitted model, macro zoobenthic carnivores were predicted to increase by 0.37% under the 10ha scenario, about double the expected increase based on the 2.4% increase for the 120ha scenario.

### 4.0. Discussion

We found that *M. pyrifera* primary productivity is an order of magnitude greater than *E. radiata* in Tasmania, indicating that *M. pyrifera* restoration at large scales could increase the primary productivity within Tasmania’s kelp forests. When this productivity benefit was incorporated into a statewide Tasmanian food web model, the biomass of many reef-associated faunal groups including commercially important species were predicted to increase. These predictions enhance the limited understanding of the ecological function of *M. pyrifera* compared to *E. radiata* in Tasmanian ecosystems (Forbes et al., 2024), and provide the first insight into the potential ecosystem benefits of *M. pyrifera* restoration in Tasmania. The predicted biomass enhancements provide expectations for how *M. pyrifera* restoration may benefit fisheries and ecosystem health. The translation of area-based restoration targets into a broader suite of expected ecosystem outcomes have the potential to boost motivation and engagement of communities and stakeholders and inform cost-benefit analyses, which are essential for successful large-scale ecosystem restoration (Jellinek et al., 2019).

The productivity ratios identified in this study are consistent with previous Tasmanian and global estimates which consistently report a greater productivity of *M. pyrifera* than *E. radiata* (Pessarrodona et al., 2022). Global comparisons of kelp species can be affected by differences in methodology and units of productivity. In addition, comparison of global estimates may misrepresent the effects of replacing one species with another at a specific site, because differences in the distribution and environmental ranges of each species mean species’ global productivity estimates are not directly comparable (Pessarrodona et al., 2019). Therefore, location-specific productivity benefits of *M. pyrifera* restoration, such as those uncovered here, are valuable for predicting ecological outcomes of restoration at specific areas.

The EwE model predicted that many reef-associated functional groups would increase in biomass following the simulation of *M. pyrifera* restoration. Predictions for the 120ha scenario were substantial given the scale of the model, whereas the 10ha and 0.7ha scenarios showed ecologically insignificant biomass and catch responses, which was expected given the changes in biomass scaled linearly with the different restoration scenarios. Biomass enhancements were predicted to translate into improved fishery catches for some key fisheries including abalone, rock lobster and some finfish. Tasmanian reef species rely on a mix of phytoplankton and macroalgae derived production (Duggins et al., 1989). The carbon fixed by kelp has various fates; it can be used by the kelp itself, lost through erosion and mortality to offshore or adjacent ecosystems (Dugan et al., 2003; Krumhansl & Scheibling, 2012), or consumed within the ecosystem via direct grazing or more likely through detrital and microbial pathways (Duggins et al., 1989; Krumhansl & Scheibling, 2012). These consumptive flows were predicted to flow through to some higher trophic level consumers including rock lobster, striped trumpeter and fur seals. Whereas groups that showed minimal changes following *M. pyrifera* restoration scenarios were likely strongly tied to a planktonic diet or limited by trophic interactions, such as increases in predator abundance (Watson et al., 2013).

The predicted direction and extent of biomass change for many functional groups following *M. pyrifera* restoration was significantly influenced by the vulnerability settings. The sensitivity of EwE model predictions to predator-prey interactions is a well-known aspect of EwE models (Heymans et al., 2016; Mackinson et al., 2009). Therefore, caution is needed when interpreting the predicted changes in biomass of functional groups following *M. pyrifera* restoration. We have higher confidence in predictions for groups that had similar directions of change under different vulnerability settings, such as was the case for many reef-associated faunal groups. We have less confidence in the predictions for groups such as shelf carnivores (e.g. flathead), small benthic carnivores and octopus, whose predictions were sensitive to the vulnerability settings. Future work is needed to estimate predator-prey interactions and ongoing monitoring of species biomass in restoration sites could help validate the predictions made here.

We suggest that the predicted biomass changes following *M. pyrifera* restoration in the model are conservative for several reasons. First, the restoration target area of 120ha is likely a lower bound estimate on the KMGBF target. The 120ha restoration target area was based on the only available statewide historical estimate of *M. pyrifera* extent, which was ∼422ha in 1997-99 (Butler et al., 2020), to align with the model scale. However, severe declines of *M. pyrifera* were recorded prior to that time and regional estimates suggest that the historical extent was likely much greater (Johnson et al., 2011). Thus, reaching the 30% target may require more than 120ha of restoration, and would result in greater predicted increases in biomass in the model simulations. This highlights a complicated aspect of applying global restoration targets into national or state scale outcomes and speaks to the importance of determining historical baselines for degraded ecosystems (Bell-James et al., 2024).

Second, Ecosim models operate at a coarse spatial resolution, representing large areas as homogenous units (Bentley et al., 2024; Heymans et al., 2016). This can mask important spatial heterogeneity and localized dynamics. For example, this study used a statewide model, assuming restoration efforts are evenly distributed across the entire reef habitat in Tasmania. However, restoration occurs at specific sites, primarily on the east coast of Tasmania. This discrepancy could lead to an underestimation of the actual impact of restoration on reef-associated species in close proximity to the restored sites and explains the ecologically insignificant predicted changes to biomass and catch following the smaller restoration scenarios. Localised ecosystem models could enhance our understanding of regional benefits of *M. pyrifera* restoration.

Lastly, the predicted biomass changes may be conservative because the model simulations did not account for the habitat benefits of kelp, which supports many animals (Forbes et al., 2024) and can enhance the recruitment of fisheries species (Hinojosa et al., 2015; Shelamoff et al., 2022). Habitat effects were excluded from the model due to a lack of understanding regarding the differences in habitat benefits of *M. pyrifera* compared to *E. radiata*. A recent study in Tasmania found that while that *M. pyrifera* forests had greater abundances of mobile fishes, there were no significant differences between invertebrates or overall community composition between the two forest types (Forbes et al., 2024). Although the two forests may serve similar ecological functions, it is possible that *M. pyrifera* forests supported unique communities before their decline. We hypothesize that *M. pyrifera* restoration would increase the availability of canopy habitat and potentially modify predator-prey dynamics within kelp forest environments. Further research on the structural roles of both *M. pyrifera* and *E. radiata* could help future models incorporate both habitat and productivity benefits when estimating vulnerability settings and simulating *M. pyrifera* restoration in Tasmania.

Several assumptions were made in this study. Here, we will discuss two key caveats.

First, Ecosim models aggregate species into functional groups based on shared characteristics, such as diet and habitat as a consequence of the uncertainty about energy flows within an ecosystem. Specifically, the flow of kelp-derived organic carbon to fisheries remains uncertain (Elliott Smith & Fox, 2022). Aggregating species into functional groups can overlook important species-specific interactions. For instance, by providing a floating surface canopy, *Macrocystis* forests modify the understorey environment differently to low lying *Ecklonia* forests which scour the seafloor. In turn understorey algal assemblages and food availability can differ substantially between forest types. Abalone in Tasmanian can modify their diet to the availability of different algal types (Shepherd, 1973), but the Tasmanian Waters EwE model did not resolve different algal species, and the flow on effects of these changes are currently unknown. Isotope analyses have identified *M. pyrifera* as an important food source for Abalone (Guest et al., 2008) and further tracing studies could uncover species-specific consumption of kelp, leading to more accurate representations of the *M. pyrifera* trophic contributions in Tasmanian food webs (Miller & Page, 2012).

A second caveat is that the model predictions did not account for future trends in climate change and fishing effort. Climate change is likely already affecting kelp forests globally as warming ocean temperatures, marine heatwaves and changes to nutrient levels have all coincided with widespread loss (Wernberg et al., 2016). Range expansions of herbivorous grazers (e.g., urchins), driven by climate change, threaten M. pyrifera restoration (Johnson et al., 2011). Further research is needed to understand how kelp forest loss from urchin overgrazing impacts fisheries productivity (Johnson et al., 2011). This study assumed that *M. pyrifera* restoration targets have been met without considering how climate change may influence restoration success. Restoration of thermally tolerant *M. pyrifera* individuals is currently underway (Layton et al., 2022) and ongoing research is investigating how climate change may affect *M. pyrifera* restoration in Tasmania (Layton & Johnson, 2021). Trajectories of climate change could be incorporated into model simulations of *M. pyrifera* restoration in the future. Fishing effort was kept constant through time however *M. pyrifera* restoration is likely to affect or be affected by changes in fishing effort. For instance, an urchin fishery has been introduced in Tasmania to manage the expansion of kelp barrens created by range-expanding urchins (NRE & FRDC, 2023). Along with the potential plans to introduce lobsters into the *M. pyrifera* restoration sites, these projected changes in fisheries will likely impact the food web dynamics in the reef system and therefore our model outputs are a simplified prediction of *M. pyrifera* restoration in a Tasmanian ecosystem.

### 4.1. Implications for conservation

Through *M. pyrifera* restoration in Tasmania, we may see improvements to the biomass of reef-associated groups as well as improvements in some fisheries. This finding has multiple implications for the way we plan and fund restoration. Kelp restoration has potential to be included in natural capital and nature repair markets that Australia is currently developing. Non-profit organisations such as OzFish Unlimited currently supports *M. pyrifera* restoration for its presumed benefits for recreational fisheries. With more evidence to suggest that *M. pyrifera* restoration will benefit fisheries species such as abalone, striped trumpeter and rock lobster, industry motivations to invest in *M. pyrifera* restoration may be improved and restoration practitioners could access a bigger pool of funding. Fisheries policy makers should further investigate the potential of restoration to assist in recovery of over or fully exploited fish stocks. Future efforts using local-scale ecosystem models and species-specific kelp groups are likely to predict greater biomass enhancements following *M. pyrifera* restoration than those predicted here, but the indications provided remain informative given their conservative nature.

